# DnaE uses strand displacement synthesis during Okazaki fragment repair

**DOI:** 10.64898/2026.04.08.717263

**Authors:** Abigail H. Kendal, Frances Caroline Lowder, Lina G. Jeffery, Lyle A. Simmons

## Abstract

RNA serves the essential role of priming DNA replication for all organisms. In bacteria, it is estimated that more than 20,000 ribonucleotides accumulate on the lagging strand per round of replication. Despite the importance of RNA primers in initiating DNA synthesis, unresolved primers result in strand breaks and induction of the DNA damage response, leading to genome instability. In bacteria, the prevailing model suggests that DNA polymerase I (Pol I) replaces RNA primers with DNA during lagging strand replication. However, in *Bacillus subtilis*, we show that Δ*polA* cells have a near wild-type phenotype, demonstrating that cells lacking Pol I still perform efficient Okazaki fragment repair. To determine how cells compensate for the loss of Pol I, we tested the ability of two major replicative polymerases, DnaE and PolC, to participate in lagging strand replication in vitro. We found that DnaE replicates an Okazaki fragment as efficiently as Pol I using strand displacement synthesis. In contrast, PolC is unable to catalyze strand displacement synthesis, but can facilitate repair when combined with a nuclease or a gapped substrate. Together, our work shows that *B. subtilis* can use several different mechanisms to replicate the lagging strand, ensuring fidelity within the replication cycle.

## INTRODUCTION

DNA replication is required for the duplication of chromosomes before they are segregated into daughter cells (1). During DNA replication, the leading strand is replicated continuously while the lagging strand is replicated in smaller pieces referred to as Okazaki fragments (2,3). Though the essential process is conserved across all life, the proteins involved in replication vary between organisms. In eukaryotic replication, synthesis of the two strands is primarily performed by DNA polymerase ε, which synthesizes the leading strand, and DNA polymerase δ, which synthesizes the lagging strand (4,5). Conversely, bacteria use a single protein complex, the DNA polymerase III holoenzyme, to replicate both the leading and lagging strands (5,6). In *Escherichia coli*, the catalytic subunit of the replicase is DnaE, which is responsible for both primer extension and bulk synthesis. The *Bacillus subtilis* protein PolC functions in an analogous role to replicate both the leading strand and the majority of the lagging strand (7,8). Unlike *Ec*DnaE, PolC is unable to extend from the 3′ hydroxyl of an RNA primer; this activity is instead performed by *Bs*DnaE (7). DnaE and PolC are C-family DNA polymerases, and both are essential for *B. subtilis* replication (6,9): DnaE extends the RNA primer before handing the substrate off to the more processive and proofreading-capable PolC (7,10,11).

For each Okazaki fragment, primase (DnaG) synthesizes short stretches of RNA to prime DNA synthesis (12,13). These primers constitute a significant source of covalently embedded genomic RNA-DNA hybrids, as it is estimated that at least 20,000 ribonucleotides are incorporated into the genome of *B. subtilis* (4.2 Mb) per round of replication exclusively due to priming activity (14). The formation of primers is essential, given that DNA cannot be synthesized de novo, yet a failure to remove RNA primers once DNA synthesis is complete results in challenges to genome integrity, such as strand breaks and increased mutation rate, or even cell death (1,15,16). In eukaryotes, mutation of proteins involved in lagging strand synthesis is linked to broad genome instability as well as the development of certain cancers (17). Thus, the maintenance of efficient lagging strand replication is imperative for genome stability in all species, ranging from bacterial to human.

The prevailing model for Okazaki fragment maturation in bacteria is that DNA polymerase I (Pol I) removes the RNA primer and replaces it with DNA, generating a nicked substrate for ligase to seal (1,22). To accomplish this, Pol I has three functional domains: an N-terminal 5′-3′ exonuclease domain, a central proofreading 3′-5′ exonuclease domain, and a C-terminal DNA polymerase domain (20,21). The combined actions of the 5′-3′ exonuclease and polymerase domains enable Pol I to simultaneously remove RNA while replacing the missing ribonucleotides with DNA (1). Other proteins have also been shown to aid Pol I in Okazaki fragment maturation. For example, in *B. subtilis*, RNase HIII cleaves the phosphodiester backbone of covalent RNA-DNA hybrids. This can shorten primers, allowing for more efficient primer removal by Pol I, thereby stimulating DNA synthesis activity (23).

Bacterial Okazaki fragment processing, as described, is based primarily on studies using the *E. coli* proteins. In one experiment, it was shown that though a deletion of *Ec*Pol I (Δ*polA)* is lethal when the cells are grown on a rich medium, these cells can grow slowly on a minimal medium, suggesting that other proteins can substitute for Pol I, albeit very poorly (24). In stark contrast to these results, a *B. subtilis* Δ*polA* strain has been reported to have only minor growth defects with sensitivity to DNA damage representing the strongest phenotype described (25,26). A key difference between the two lineages is that *B. subtilis*, like many gram-positive bacteria, encodes a second discrete 5′ to 3′ nuclease referred to as FEN or FenA (25-28). It has been shown that *B. subtilis* FEN exhibits 5′-3′ exonuclease activity and substrate-specific flap endonuclease activity, though, like Pol I, deletion of FEN (Δ*fenA*) does not cause a growth defect (25). A deletion of both *polA* and *fenA*, however, is synthetically lethal (26). This demonstrates that the 5′-3′ exonuclease activity is essential in *B. subtilis*, though DNA synthesis can be provided by a protein other than Pol I. The identity of this other polymerase that enables Pol I-independent Okazaki fragment maturation has yet to be discovered.

In this study, we explore the consequences of impaired Okazaki fragment processing on *B. subtilis* cell growth and viability. We found that during normal growth in rich medium, loss of Pol I activity had only a very minor effect on cell length with no effect on chromosome partitioning or induction of the SOS response to DNA damage. Cells further compromised in lagging strand repair exhibit significant cell elongation, chromosome partitioning defects, and induction of the SOS response. To understand how cells replicate the lagging strand in the absence of Pol I, we tested purified PolC and DnaE on model Okazaki fragment substrates in vitro. We found that DnaE catalyzes robust strand displacement synthesis from a gap, while PolC arrests at the 5′ end of the downstream Okazaki fragment in the absence of a nuclease. Further, we found that DnaE works in conjunction with FEN to more efficiently process the lagging strand, likely representing the major pathway for Okazaki fragment repair in the absence of Pol I. In the presence of a nuclease that can sufficiently degrade a downstream fragment, we report that PolC is capable of repair, albeit less efficiently, suggesting that PolC-directed Okazaki fragment processing represents a minor repair pathway. Together, our work shows that DnaE and FEN are able to efficiently compensate for the loss of Pol I activity. Importantly, our work also shows that bacteria contain multiple pathways for Okazaki fragment processing, ensuring fidelity in the overall replication process.

## MATERIALS AND METHODS

### Bacteriology

All strains used were derived from *Bacillus subtilis* PY79 using standard procedures as described (29,30)**(Supplementary Table 1)**. Cells were grown in lysogeny broth (LB)(31). Antibiotics were used at the following concentrations: spectinomycin (100 µg/mL), kanamycin (50 µg/mL), and chloramphenicol (5 µg/mL).

### Spot titers

All strains were stuck onto LB agar plates with the appropriate antibiotic for overnight growth at 37°C. The following morning, LB supplemented with the appropriate antibiotic was inoculated with a single colony and grown at 37°C in a rolling rack until an OD_600_ between 0.5 and 0.7 was reached. Samples were normalized to an OD_600_ of 0.5 using 0.85% saline and serial dilutions were performed in a 96-well plate. An equivalent volume of each dilution was spotted onto LB agar plates with no drug treatment, 20 ng/mL mitomycin C (MMC), 100 µg/mL methyl methane sulfonate (MMS), 20 ng/mL ciprofloxacin, or 5 mM hydroxyurea (HU) and grown at 37°C for 12 hours. Whole image contrast was adjusted in FIJI.

### Microscopy

Each strain was struck onto LB agar plates and grown overnight at 30°C. The next morning, S7_50_ minimal media (1x S7(50) [10x S7(50): 0.5 M MOPS, 100 mM (NH_4_)2SO_4_, 50 nM KH_2_PO_4_], 1x metals [100x metals: 0.2 M MgCl_2_, 70 mM CaCl_2_, 5 mM MnCl_2_, 0.1 mM ZnCl_2_, 100 µg/mL Thiamine-HCl, 2 mM HCl, 0.5 mM FeCl_3_,], 1% glucose, 0.1% glutamate) was used to wash the plates for a starting OD_600_ of 0.15 in S7_50_ as described (32-34). To avoid light-induced changes in the media, the liquid culture was protected from light by covering the flask with aluminum foil. The flask was incubated in a shaking water bath at 30°C with 250 rpm. The OD_600_ was measured until cells reached exponential phase (OD_600_ = 0.6-0.8). Each culture was mixed with the appropriate dye. For membrane staining, FM4-64 dye was added to a final concentration of 3.33 µg/mL. For the nucleoid staining, 4′,6-diamidino-2-phenylindole (DAPI) was added to 1.5 µg/mL. Cells were then spotted onto 15-well microscope slides with 1% agarose pads in 1X S7_50_ salts. Slides were left at room temperature for 5-10 minutes before aspirating residual fluid and attaching a coverslip. An Olympus BX61 fluorescence microscope and the Olympus Cellsens software were used to capture images of cells as described (32-34). For each experiment, several hundred cells from each strain were imaged over two different days using the same growth conditions.

### Image analysis

Images were analyzed using ImageJ. To assess nucleoids, corresponding images with membrane stain and DAPI were overlayed using the Olympus Cellsens software. Fluorescence intensity thresholds were set to determine if a cell was undergoing SOS induction using the Microbe plugin in FIJI. Cells with fluorescence per area above 1.2 were considered positive for SOS induction (32).

### Statistical analysis

Analysis of cell and nucleoid length data was completed using base R. Data were assessed for normalcy using a Shapiro-Wilk test to calculate a W statistic with a corresponding p-value. A Kruskal-Wallis rank sum test was used as a non-parametric multiple comparison analysis tool. For data sets with a significant difference, a Mann-Whitney U test (Wilcoxon rank sum test) with continuity correction was used to evaluate individual relationships. For each test of significance performed, α = 0.05. The *n* associated with each test can be found in the figure legends. Significance is denoted in figures as follows: * = p < 0.05, ** = p < 0.01, *** = p < 0.001.

### Genomic DNA purification

LB inoculated with the appropriate *B. subtilis* strain was grown to OD_600_ ∼ 0.7. The cells were pelleted and resuspended in lysis buffer (10 mM Tris-HCl pH 8, 10 mM EDTA pH 8, 1% Triton X-100, 0.5 mg/mL RNase A, and 20 mg/mL lysozyme), then incubated at 37°C for 30 minutes. SDS (1% final concentration) and protease K (final concentration of 1.3 mg/mL) were added, followed by incubation at 55°C for 30 minutes. The volume was loaded onto a silica spin column and a 2x volume of PB buffer was added (5M guanidine-HCl, 30% isopropanol). Samples were centrifuged at 12,000 x g for 1 minute. PB buffer was used to wash the column followed by centrifugation at 12,000 x g for 1 minute. The column was washed with PE buffer (10 mM Tris-HCl pH 7.5, 80% ethanol) and centrifuged at 12,000 x g for 1 minute. To dry the column, centrifugation at 14,000 x g was performed for 3 minutes. The column was eluted with nuclease-free water and centrifuged at 14,000 x g for 1 minute. The eluate was reloaded onto the column and centrifuged at 14,000 x g for 1 minute.

### Cell competency and transformation

A single colony was inoculated into LM media (LB supplemented with 3mM MgSO_4_) and grown until an OD_600_ of 0.8 was reached. This culture was used to inoculate MD media (1x PC buffer, 2% glucose, 11 µg/mL ferric ammonium citrate, 2.5 mg/mL potassium aspartate, 3 mM MgSO_4_) which was shaken at 37°C for 4 hours. These competent cells were transformed with 100 ng of genomic DNA and incubated on a rolling rack at 37°C for 90 minutes. The entire volume was plated on LB agar plates with the appropriate antibiotic then incubated overnight at 37°C.

### Protein expression

Chemically competent *E. coli* BL21(DE3) were transformed with the appropriate plasmid **(Supplementary Table 4)** and plated on LB agar plates supplemented with kanamycin (Kan, final concentration of 25 µg/mL) and grown overnight at 37°C. These colonies were used to inoculate an LB + Kan starter culture, which was grown overnight at 37°C while shaking. After inoculation with overnight culture, each liter of LB + Kan was grown to an OD_600_ ∼ 0.7, at which point cells were induced by adding isopropyl β-D-1-thiogalactopyranoside (IPTG) to 0.5 mM. Cells expressing RNase HIII were grown for 3 hours at 37°C while cells expressing FEN or Pol I were grown for 18 hours at 25°C. In both cases, the cells were harvested by centrifugation at 4000 x g/4°C/25 minutes. After discarding the supernatant, cell pellets were stored at -80°C.

### Protein purification

After thawing on ice, protein pellets were resuspended in lysis buffer (20 mM Tris pH 8.0, 400 mM NaCl, 5% v/v glycerol), supplemented with EDTA-free protease inhibitor (Pierce: A32965). The cells were lysed via sonication, and the lysate was clarified by centrifugation at 30000 x g for 30 minutes. The clarified lysate was applied to a Ni^2+^-NTA-agarose gravity column, which was equilibrated with lysis buffer. The column was washed with 5 CV lysis buffer and 5 CV wash buffer (20 mM Tris pH 8.0, 2 M NaCl, 15 mM imidazole, 5% v/v glycerol), then the bound protein was eluted using 3 CV elution buffer (20 mM Tris pH 8.0, 400 mM NaCl, 300 mM imidazole, 5% v/v glycerol). Protein containing fractions were pooled and diluted such that the final NaCl concentration was 240 mM then treated with Ulp-1 protease for 2 hours at 25°C. The cleaved protein was dialyzed overnight against dialysis buffer (20 mM Tris pH 8.0, 300 mM NaCl, 5% v/v glycerol). The dialyzed protein was passed over a Ni^2+^-NTA-agarose gravity column equilibrated with dialysis buffer and the flowthrough was collected. The column was next washed with 3 CV dialysis buffer, with the flowthrough collected. These fractions were pooled and diluted to 50 mM NaCl with Q buffer A (20 mM Tris pH 8.0, 5% v/v glycerol, 1 mM DTT). The diluted protein was filtered and applied to an ion exchange column as appropriate. A HiTrap Q FF anion exchange column (Cytiva, 17515601) was used for FEN and Pol I while a HiTrap SP HP cation exchange column (Cytiva, 17115101) was used for RNase HIII and fractionated with an AKTA FPLC. The protein was eluted using an increasing gradient of Q buffer B (20 mM Tris pH 8.0, 500 mM NaCl, 5% v/v glycerol, 1 mM DTT), then peak fractions were evaluated using SDS-PAGE. After pooling and concentrating the protein using a centrifugal filter, glycerol was added to 25%. Then the proteins were aliquoted, flash-frozen, and stored at -80°C. The gel of proteins used in this study was made by separating 2 µg of each protein using a Mini-PROTEAN TGX 4-20% gradient gel (BIORAD, 4561096) then staining with Coomassie blue **(Supplementary Figure 1)**. This purification method was used for FEN, Pol I, and RNase HIII. We were generously given purified PolC and DnaE by Charles McHenry.

### Okazaki fragment repair assays

To create the substrate, oligonucleotides were mixed to 1000 nM in 1x dilution buffer (20 mM Tris pH 8, 50 mM NaCl) and annealed by heating at 98°C for 1 minute followed by cooling in the dark for 30 minutes. All oligonucleotides were purchased from IDT and sequences are included in **Supplementary Table 2**. Substrate compositions are provided in **Supplementary Table 3**. Pure proteins were diluted to 400 nM using 1x extension reaction buffer (80 mM Tris-acetate pH 7.8, 24 mM magnesium acetate, 600 mM potassium glutamate, 6 µM zinc sulfate, 4% wt/vol polyethylene glycol 8000, 0.04% Pluronic F68) and added to a reaction mix of 50 µM dNTPs, 100 nM substrate, 1x extension reaction buffer, and nuclease-free water. The final polymerase concentration for each reaction was 100 nM. Reactions were performed at room temperature for 45 minutes, with an optional 1 minute pre-incubation with FEN or RNase HIII (40 nM final concentration unless otherwise noted in the figure legend). Reactions were quenched using an equal volume of reaction stop buffer (95% formamide, 20 mM EDTA, 0.01% bromophenol blue) and boiled at 98°C for 10 minutes followed by snap cooling on ice. To ensure full denaturation of nucleic acid-protein complexes, 1x TBE running buffer was heated to 50°C prior to electrophoresis. Assays were visualized using urea-PAGE with 20% acrylamide and 8M urea and electrophoresis was performed on a hot plate set to 50°C. Gels were visualized using the 700 and 800 nM channels of a LiCor Oddesey CLx. LUT colors were altered from red/green to magenta/green in FIJI for accessibility purposes.

Relative extension products were quantified from the gels by isolating the channel for the labeled primer and taking the area under the curves for each lane. Partial extension was considered any peak between the initial primer and full-length product, which can both be identified using the included ladder. Any bands lower than the initial primer were considered degradation products. The percentage of each outcome was calculated with respect to the total signal for the lane. A Kruskal-Wallis test was used to assess for significant differences across the treatment groups, and the effect of the added nuclease on the fraction of each type of extension. Pairwise Wilcoxon rank-sum tests were performed with Benjamini-Hochberg correction for multiple comparisons of each nuclease + polymerase lane to the relevant polymerase-only lane. Significant pairwise comparisons (p.adj < 0.05) were annotated using the following: * = p < 0.05, ** = p < 0.01, *** = p < 0.001.

### Tecan fluorescence reporter assay

The indicated strains were struck onto LB agar plates for overnight growth at 30°C. These cells were used to inoculate LB cultures which were grown until an OD_600_ between 0.75 and 1.25 was reached. Strains were diluted to OD_600_ values of 0.25, 0.5, and 0.75 using 0.85% saline in a black 96-well plate with a clear bottom panel. Relative fluorescence was measured using a customized method optimized for the mCitrine fluorophore in a Tecan Spark plate-reader. Three biological replicates were performed, and each biological replicate consisted of two technical replicates.

A linear mixed-effects model was fit with sample identity and OD as fixed effects, including their interaction, and biological replicate as a random intercept. Estimated marginal means were extracted from the fitted model and pairwise contrasts between each deletion strain and wild-type were computed separately at each OD level using the emmeans package in R. P-values were adjusted for multiple comparisons using the Benjamini-Hochberg method, and comparisons with adjusted p < 0.05 were considered statistically significant. Non-significant comparisons are omitted from the graph and significant comparisons are indicated as follows: * = p < 0.05, ** = p < 0.01, *** = p < 0.001. Error bars indicate ±1 SD.

## RESULTS

### Double deletion mutants in lagging strand replication cause the most severe growth interference when challenged with DNA damage

Since *B. subtilis* cells without DNA polymerase I show a near wild-type (WT) phenotype during normal growth, we asked how other mutants compromised for DNA replication would respond when challenged with low levels of DNA damage **(Figure 1A)**. Cells with defects in Okazaki fragment maturation are sensitive to exogenous DNA damage, providing a proxy for assessing protein involvement during lagging strand synthesis (24,25). Therefore, we performed spot dilution assays of WT, Δ*polA*, Δ*rnhC*, Δ*fenA*, Δ*rnhC*Δ*fenA*, and Δ*rnhC*Δ*polA* cells to compare their sensitivity to low levels of DNA damage induced by treatment with hydroxyurea (ribonucleotide reductase inhibitor), mitomycin C [MMC] (alkylating agent), methyl methane sulfonate [MMS] (alkylating agent), and ciprofloxacin (fluoroquinolone). We show that strains lacking one protein (Δ*polA*, Δ*rnhC*, or *ΔfenA*) have a mildly sensitive phenotype, while the double deletion strains Δ*rnhC*Δ*fenA* and Δ*rnhC*Δ*polA* are hypersensitive to each type of DNA damage tested. Thus, deletion of one component of the canonical Okazaki fragment processing pathway imparts only a minor sensitivity to DNA damage, whereas inactivating two components has a much stronger effect. These data suggest that *B. subtilis* cells use multiple pathways for Okazaki fragment processing to help mitigate the effects of DNA damage that would otherwise impact DNA replication, increase mutation rate, and potentially lead to cell death.

**Figure 1.**
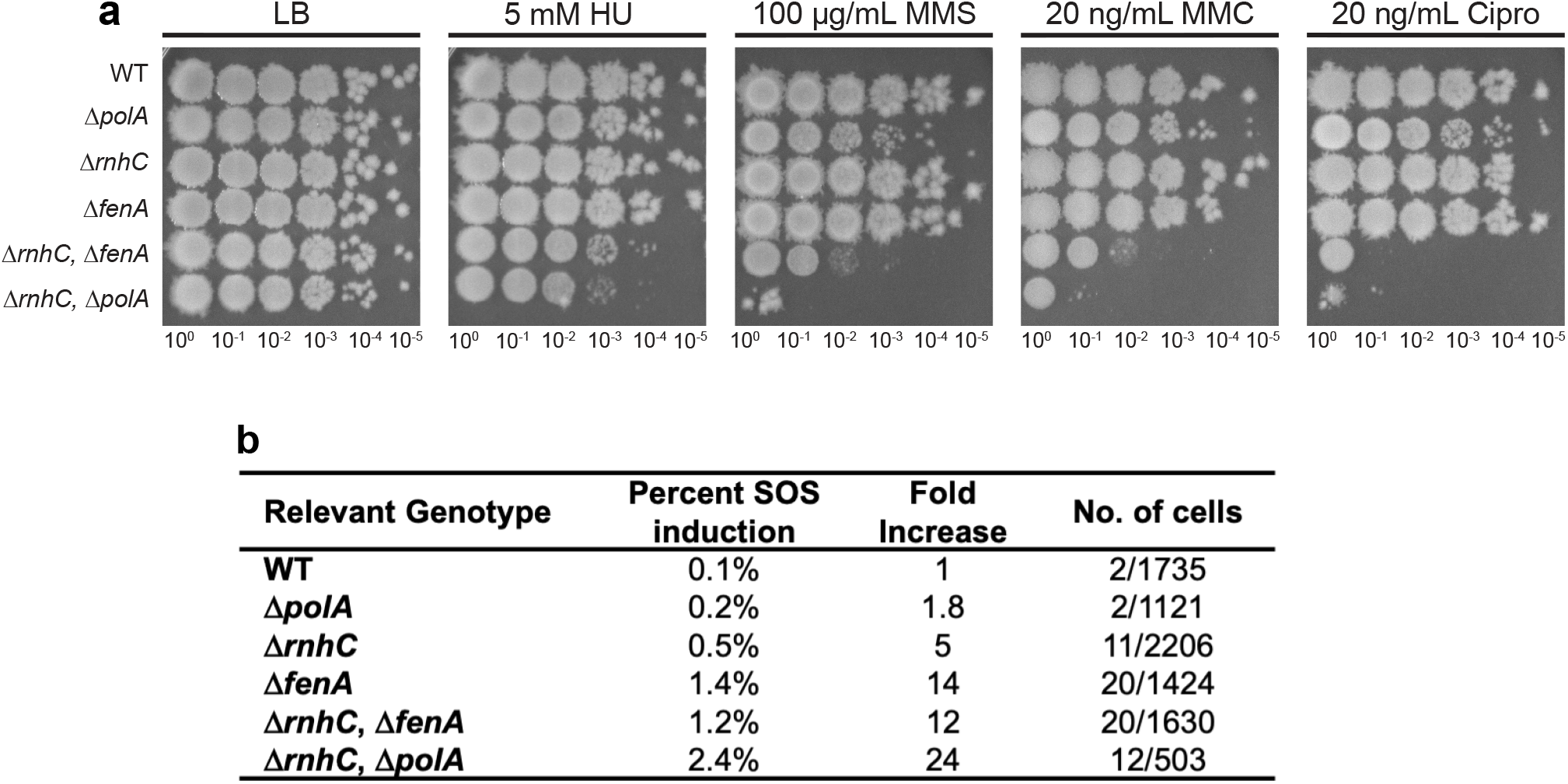
Defects in lagging strand processing result in anucleate cells, guillotined chromosomes, induction of the SOS response, and sensitivity to DNA damaging agents. **(A)** Spot titer assay with increasing serial dilutions of the indicated strains. Strains were plated on LB medium with the indicated concentration of DNA damaging agent: hydroxyurea (HU), methylmethane sulfonate (MMS), mitomycin C (MMC) or ciprofloxacin (Cipro). Spot titer growth was verified through the completion of three biological replicates. **(B)** SOS induction was measured by DinC-GFP reporter foci for the genotypes listed using live-cell fluorescence microscopy. The number of cells scored and the number positive for SOS induction are indicated.

### Compromised lagging strand replication result in SOS induction

The hallmark of a *Ec*Pol I deletion (Δ*polA)* is induction of the DNA damage response (SOS) (35). In the absence of Pol I, current *E. coli* literature suggests that excessive nicks and gaps result from failures in Okazaki fragment maturation, causing hyper-recombination and induction of the SOS response (35). To measure SOS induction at the single-cell level, we fused GFP to an SOS reporter gene known as *dinC*, one of the most highly expressed SOS genes in *B. subtilis* (36). The *dinC-gfp* allele was expressed at its native locus in WT, Δ*polA*, Δ*rnhC*, Δ*fenA*, Δ*rnhC*Δ*fenA*, and Δ*rnhC*Δ*polA* under normal growth conditions to assay for SOS induction as another indicator for compromised lagging strand replication. We found that Δ*polA* cells were only 1.8-fold induced relative to WT **(Figure 1B)**, demonstrating that the loss of Pol I has a minor effect on lagging strand replication in *B. subtilis*, as cells do not show the consequences expected if Okazaki fragment processing was severely impaired. As a control, we show that Δ*rnhC* cells were 14-fold induced for SOS, which is supported by prior work (37). The SOS induction measurements in double mutants show that Δ*rnhC*Δ*polA* has the most significant effect, causing a 24-fold increase in SOS induction relative to WT. We conclude that the loss of Pol I activity is largely benign, suggesting that a different DNA polymerase compensates in its absence.

### Defects in Okazaki fragment processing cause chromosome segregation defects and cell filamentation

To better understand the effect of defects in Okazaki fragment processing on chromosome biology, we scored the frequency of atypical nucleoid morphologies, including anucleate cells and guillotined nucleoids to single-cell resolution. We assessed a variety of mutant strains by fluorescence microscopy, scoring over 400 cells per strain. This was accomplished by growing WT, Δ*polA*, Δ*rnhC*, Δ*fenA*, Δ*rnhC*Δ*fenA*, and Δ*rnhC*Δ*polA* to mid-exponential phase followed by fluorescence microscopy with a membrane stain (FM4-64) and the DNA stain DAPI (4′,6-diamidino-2-phenylindole) for imaging. We found that WT, Δ*polA*, Δ*fenA* cells were very similar with regard to nucleoid number and chromosome segregation. Cells with a Δ*fenA* genotype did show a higher percentage of cells with one nucleoid and a lower percentage of cells with two nucleoids, suggesting an increase in crossovers between sister chromosomes **(Figure 2D)**. In contrast, cells that are Δ*rnhC*, Δ*rnhC*Δ*fenA*, and Δ*rnhC*Δ*polA* show guillotined chromosomes, anucleate cells, cell elongation, and large cytosolic space between the cell membrane and the nucleoid **(Figure 2A)**. We measured cell length and nucleoid length along the long axis of the cell or the nucleoid **(Figure 2B)**. We found that Δ*polA* cells are shorter than WT, while Δ*fenA* cells are not significantly different. Cells that are Δ*rnhC*, Δ*rnhC*Δ*fenA*, and Δ*rnhC*Δ*polA* are significantly longer than WT, supporting the phenotypes observed in the cell images. Measurement of nucleoid length yields similar results **(Figure 2C)**. Cells that are Δ*fenA* or Δ*polA* were the same as WT with respect to nucleoid length, while Δ*rnhC*, Δ*rnhC*Δ*fenA*, and Δ*rnhC*Δ*polA* cells showed abnormally long nucleoids. The long nucleoids suggest that hyper recombination may have occurred, possibly caused by nicks and gaps in the DNA resulting from problems with Okazaki fragment maturation. The observed number of anucleate, multinucleate, and guillotined nucleoid cells is summarized in **Figure 2D**. Further, these results suggest that loss of Pol I is again compensated for by the presence of other polymerases since minimal changes in cell length and nucleoid length are observed in Δ*polA* cells.

**Figure 2.**
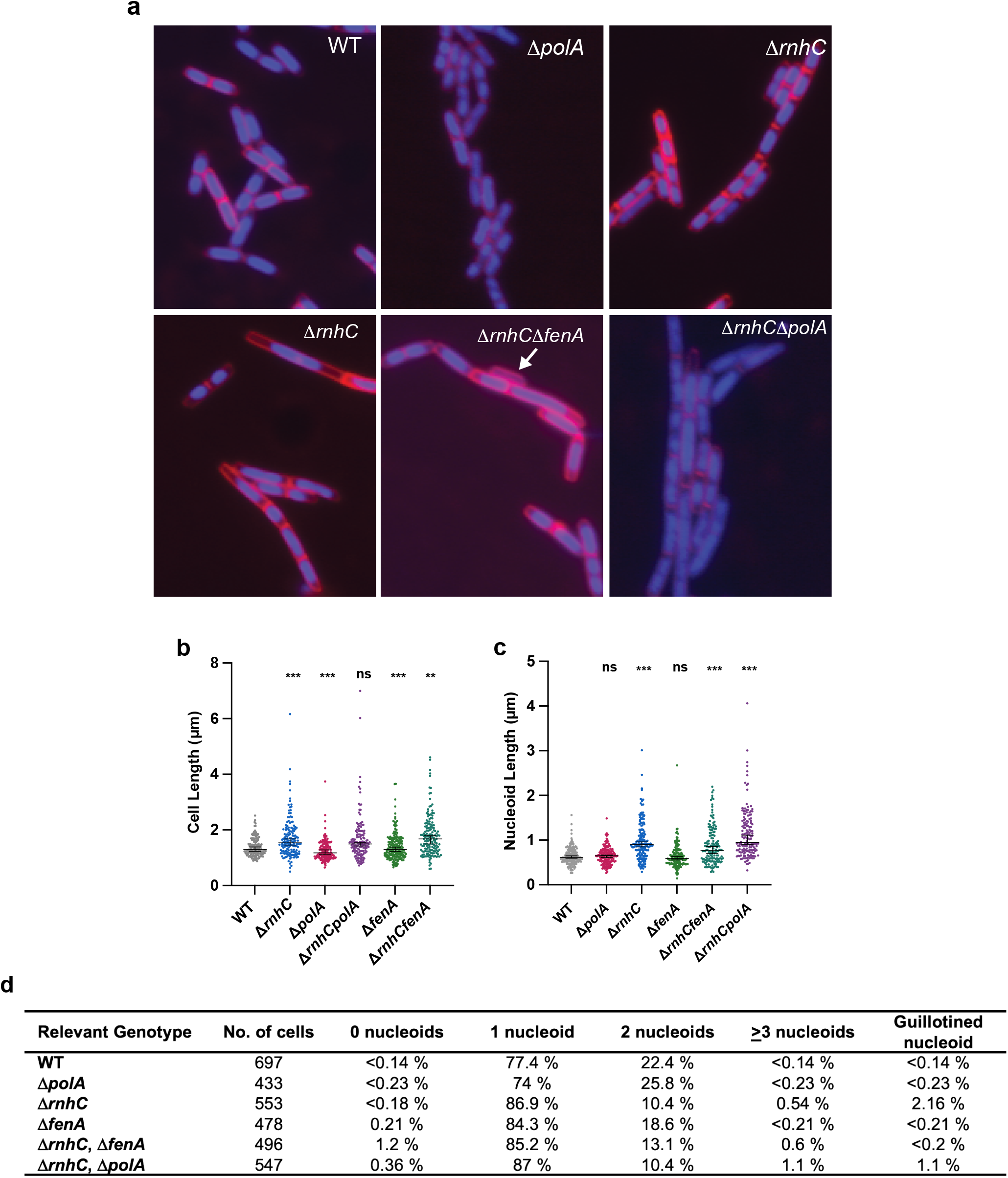
Defects in Okazaki fragment repair can be visualized through live-cell fluorescence microscopy. **(A)** Live cell fluorescence microscopy images of the strains indicated. Cell membranes are stained with the dye FM4-64 and nucleoids are visualized through DAPI staining. The white arrow indicates an anucleate cell. **(B)** Cell length measurements of the indicated strains following live-cell microscopy. Number of cells scored for each strain is noted in (D). **(C)** Nucleoid length measurements of the indicated strains following live-cell microscopy. Number of cells scored for each strain is noted in (D). **(D)** Following live-cell fluorescence microscopy, each nucleoid was scored and sorted into the following categories: 0, 1, 2, ≥3, based on the number of visible nucleoids per cell. A nucleoid was considered guillotined if a complete septum was visible over the DNA. Number of cells scored is included in the figure.

### DnaE is efficient at strand displacement synthesis from a gap

It was recently shown that Pol I has weak 5′ to 3′ exonuclease activity and the main contribution of Pol I to Okazaki fragment maturation in *B. subtilis* comes from its DNA polymerase activity (25). Additionally, our work shows that Pol I can be deleted with little phenotype, indicating that at least one other DNA polymerase can efficiently compensate for the loss of Pol I. Therefore, we tested the other major DNA polymerase(s) to determine if either or both can replace Pol I activity in vitro. Pol I is an A-family DNA polymerase and is the only A-family polymerase in *B. subtilis* (38). PolC and DnaE are C-family, major replicative DNA polymerases, and both lack 5′ to 3′ exonuclease or flap endonuclease activity (9). Since *B. subtilis* Pol I has weak 5′ to 3′ exonuclease, it relies on stand displacement synthesis to perform Okazaki fragment maturation (23,25). We reasoned that PolC or DnaE might be capable of strand displacement activity, enabling Okazaki fragment maturation in the absence of Pol I. We tested Pol I, PolC, and DnaE for activity using a DNA primer extension assay as a control on a substrate lacking a downstream fragment **(Figure 3A)**. We demonstrate that each DNA polymerase is active under the assay conditions, as all were able to fully replicate the substrate. Next, we asked whether PolC or DnaE were able to synthesize using a nicked substrate, which would require strand displacement activity. We found that Pol I can replicate DNA despite the close proximity of the downstream fragment, though PolC and DnaE were unable to extend from the nick, supporting prior observations **(Figure 3B)** (23,39). However, inside cells, gaps are expected between adjacent Okazaki fragments (40-42). Thus, we designed a 10-nucleotide gapped substrate to more accurately mimic an in vivo Okazaki fragment. With this substrate, DnaE catalyzed strand displacement synthesis and generated full-length products like Pol I, while PolC synthesis arrested at the 5′ end of the downstream fragment **(Figure 3C)**. DnaE has previously been shown to catalyze strand displacement synthesis with an existing flap (43), however, this activity has not been tested in a gapped substrate. During the Okazaki fragment maturation process, DnaE is unlikely to encounter a downstream flap and is instead more likely to encounter an annealed 5′ end with a gap between fragments, as shown in **Figure 3C**. Therefore, our results demonstrate that DnaE is capable of stand displacement synthesis when encountering an annealed 5′ end, provided there is a sufficient gap between the upstream and downstream DNA. With this data, we suggest that DnaE can process Okazaki fragments in vivo.

**Figure 3.**
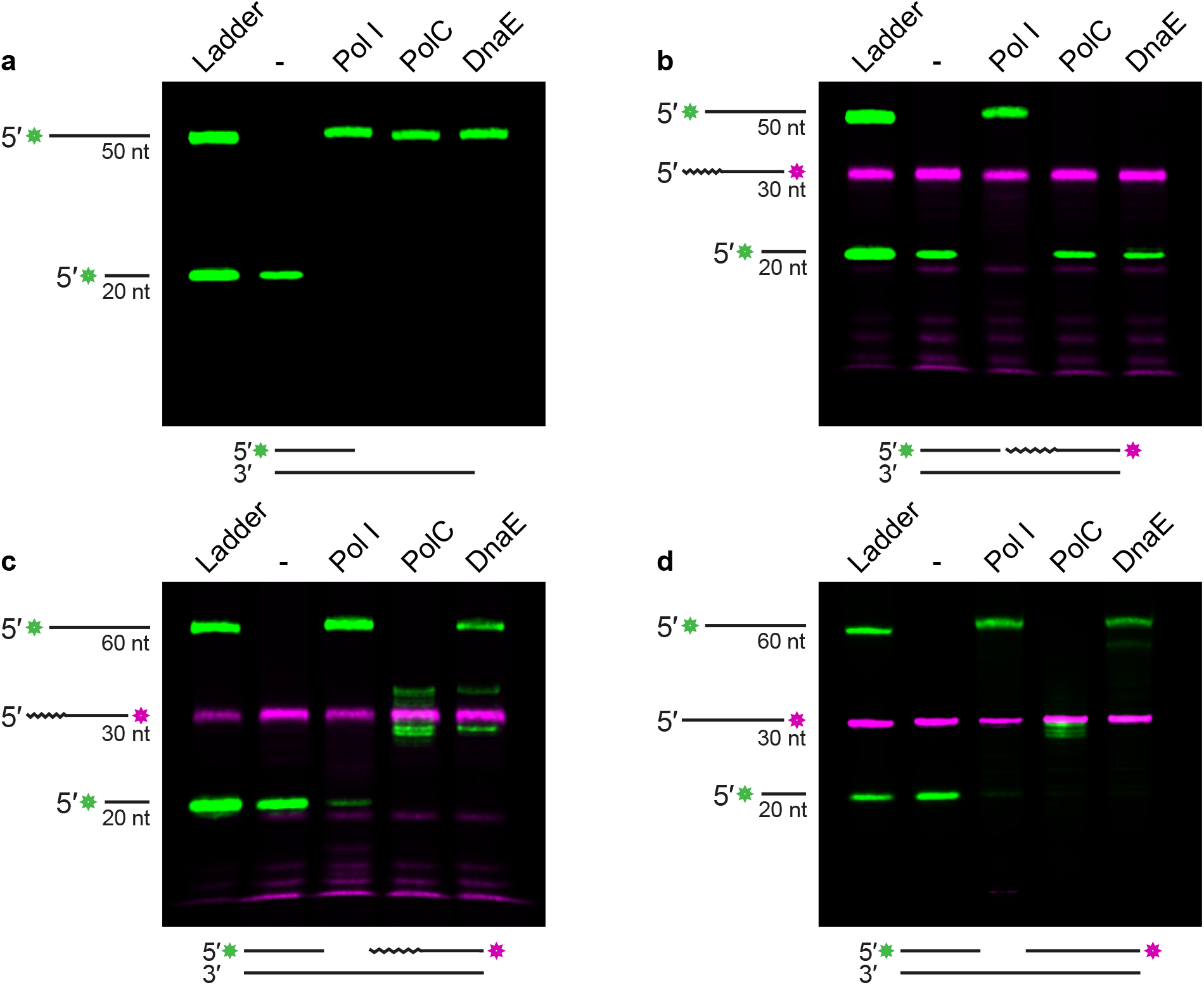
DnaE performs strand displacement synthesis on an Okazaki-fragment like gapped substrate. **(A)** Products of an extension assay using a primed substrate visualized by urea-PAGE. Ladder shows length of primer and full-length extension product. **(B)** Products of an extension assay using a nicked substrate visualized by urea-PAGE. Ladder shows length of primer, downstream fragment, and full-length extension product. Embedded ribonucleotides denoted by squiggly line. **(C)** Products of an extension assay using a 10 nt gap substrate visualized by urea-PAGE. Ladder shows length of primer, downstream fragment, and full-length extension product. Embedded ribonucleotides denoted by squiggly line. **(D)** Products of an extension assay using a DNA-only 10 nt gap substrate visualized by urea-PAGE. Ladder shows length of primer, downstream fragment, and full-length extension product.

### DnaE can perform strand displacement synthesis on a DNA-only substrate

Next, we asked if the strand displacement synthesis ability of DnaE is specific to a substrate with embedded ribonucleotides (RNA-DNA hybrid) or if strand displacement can occur with DNA. We did so using a version of the gapped substrate where the downstream strand consisted entirely of DNA. As shown in **Figure 3D**, DnaE can replicate through the downstream DNA fragment, resulting in a full-length extension product. Additionally, we see robust strand displacement activity from Pol I and minimal activity from PolC using this substrate, with no full-length extension product formed. Thus, for the proteins tested in this assay, there are minimal differences between a substrate with incorporated ribonucleotides and a DNA-only substrate. These results are supported by work with *Streptococcus pyogenes*, which suggests that DnaE may function in DNA repair and lesion bypass when a replication fork stalls (44). Consistent with existing literature in other gram-positive bacteria, our data suggest that DnaE may aid in other forms of DNA repair beyond Okazaki fragment maturation where DNA strand displacement synthesis is important.

### DnaE removes Okazaki fragments with help from FEN

Once we established that DnaE provides robust strand displacement synthesis activity, we asked if RNase HIII or FEN can enhance DnaE activity, allowing for more full-length extension. To test this, we pre-incubated the model Okazaki fragment system with RNase HIII or FEN before performing an incubation with Pol I, PolC, or DnaE and monitoring extension through the downstream Okazaki fragment **(Figure 4A)**. We found that Pol I was able to complete replication of the model substrate with or without the addition of RNase HIII or FEN, with the majority of the total signal representing full-length extension product **(Figure 4B)**. Thus, Pol I activity on the model substrate is not enhanced with the addition of either protein on a 30 nt-long substrate **(Figure 4C)**. This is likely due to the strand displacement synthesis performed by Pol I, rather than its nuclease activity, since minimal degradation of the downstream fragment was observed **(Figure 4A, 4B)**. However, we observed a sizable shift in the gel when Pol I was combined with a high concentration of FEN, indicating that FEN and Pol I interact on the substrate, suggesting these proteins may also interact in vivo **(Figure S2B)**. PolC showed no full-length product on its own or in the presence of RNase HIII. When FEN was added, PolC replication activity increased, and some full-length product was detected. We also detected smaller products, suggesting that when PolC reaches the end of the template, it uses its 3′ to 5′ exonuclease “proofreading” activity to degrade a portion of the full-length product **(Figure 4B)**.

**Figure 4.**
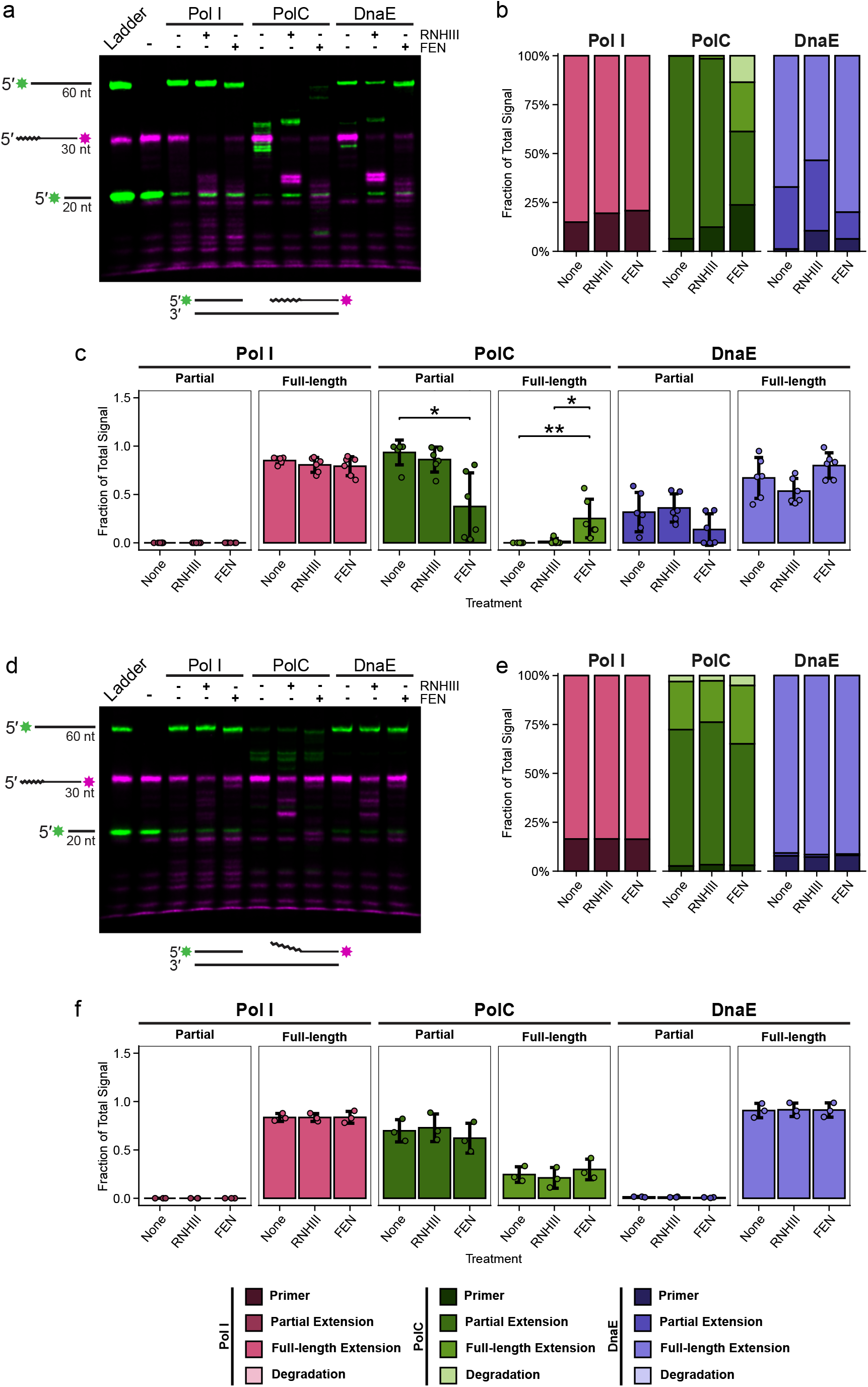
Strand displacement synthesis is improved by degradation of downstream Okazaki fragment. **(A)** Products of an extension assay using a 10 nt gap substrate visualized by urea-PAGE with the indicated proteins. Ladder shows length of primer, downstream fragment, and full-length extension product. Embedded ribonucleotides denoted by squiggly line. **(B)** Analysis of total gel signal, subsetted by primer, partial extension, full-length extension, or degradation. Color legend included below. Quantification is representative of six gel replicates, with mean values shown on the graph. **(C)** Analysis of partial and full-length extension achieved by polymerases with nuclease treatment. Quantification is representative of six gel replicates, with individual data points shown and ±1 SD indicated by the error bars. **(D)** Products of an extension assay using a 10 nt gap with flap substrate visualized by urea-PAGE with the indicated proteins. Ladder shows length of primer, downstream fragment, and full-length extension product. Embedded ribonucleotides denoted by squiggly line. **(E)** Analysis of total gel signal by primer, partial extension, full-length extension, and degradation. Color legend included below. Quantification is representative of three gel replicates, with mean values shown on the graph. **(F)** Analysis of partial and full-length extension achieved by polymerases with nuclease treatment. Quantification is representative of three gel replicates, with individual data points shown and ±1 SD indicated by the error bars.

We found that the ability of PolC to produce a full-length product using a gapped substrate is dependent on the concentration of FEN added, where a higher concentration of FEN leads to more full-length extension **(Figure S2C)**. Like Pol I, DnaE performed extension on its own, although we did detect an increase in full-length product formation when FEN was added **(Figure 4B)**. We quantified extension and found that the addition of FEN results in an increase in full-length product formation for PolC and DnaE, with the difference being statistically significant for PolC **(Figure 4C)**. With these results, we suggest that DnaE and FEN provide the best pathway for Okazaki fragment maturation in the absence of Pol I, given the robust strand displacement activity exhibited by DnaE and its enhancement in the presence of FEN.

### Downstream flaps stimulate strand displacement synthesis

Prior work showed that DnaE is able to displace a strand with a 5′ flap (43). Therefore, we asked if Pol I or PolC activity is also enhanced by a downstream fragment with a 5′ flap. If so, it would suggest that PolC could exchange for DnaE or Pol I once the flap is created, since the 5′ end of the downstream fragment would no longer be annealed to the template. We show that Pol I is able to replicate through the modeled Okazaki fragment with a flap **(Figure 4D)**. Addition of RNase HIII or FEN did not stimulate Pol I activity as expected since these proteins do not stimulate Pol I when the 5′ end is annealed **(Figure 4E, 4F)**. When PolC was tested, only weak strand displacement activity was detected, and the addition of FEN or RNase HIII did not enhance PolC activity or the accumulation of partial or full-length extension products **(Figure 4D - 4F)**. Similar to Pol I, DnaE showed robust extension with and without RNase HIII or FEN, and the amount of full-length product detected did not change significantly with the addition of RNase HIII or FEN **(Figure 4D - 4F)**. However, the addition of the flap to the substrate increased the fraction of full-length product relative to the gapped substrate for both PolC and DnaE **(Figure 4B, 4E)**. Therefore, a downstream flap effectively substitutes for FEN and enhances DnaE strand displacement activity and allows PolC to achieve weak full-length extension. With these results, we suggest that Pol I and DnaE displace a downstream Okazaki fragment-like substrate and that weak full-length extension by PolC is dependent on the presence of a nuclease or a flapped substrate **(Figure 4A, 4D)**.

### DnaE expression increases in the absence of other replication proteins

Our current results show that DnaE is capable of substituting for Pol I during lagging strand replication in vitro. Therefore, we asked if basal expression of DnaE is sufficient to perform Okazaki fragment repair in the absence of FEN, Pol I, or RNase HIII, or if DnaE expression increases when lagging strand replication is compromised. Using a fluorescence reporter assay to measure DnaE-mCitrine abundance, we found that DnaE expression does increase in the absence of FEN, Pol I, or RNase HIII **(Figure 5)**. While the magnitude of these differences varies slightly depending on the optical density of the culture, differences in relative fluorescent units (RFU) between WT DnaE-mCitrine, Δ*rnhC*, and *ΔpolA* were consistent and statistically significant, reflecting an upregulation in DnaE expression in genetic backgrounds compromised for lagging strand replication. With these results, we suggest that DnaE expression increases to circumvent the loss of Pol I or defects in nucleases involved in lagging strand replication, aiding in the ability of DnaE to engage in this process in vivo.

**Figure 5.**
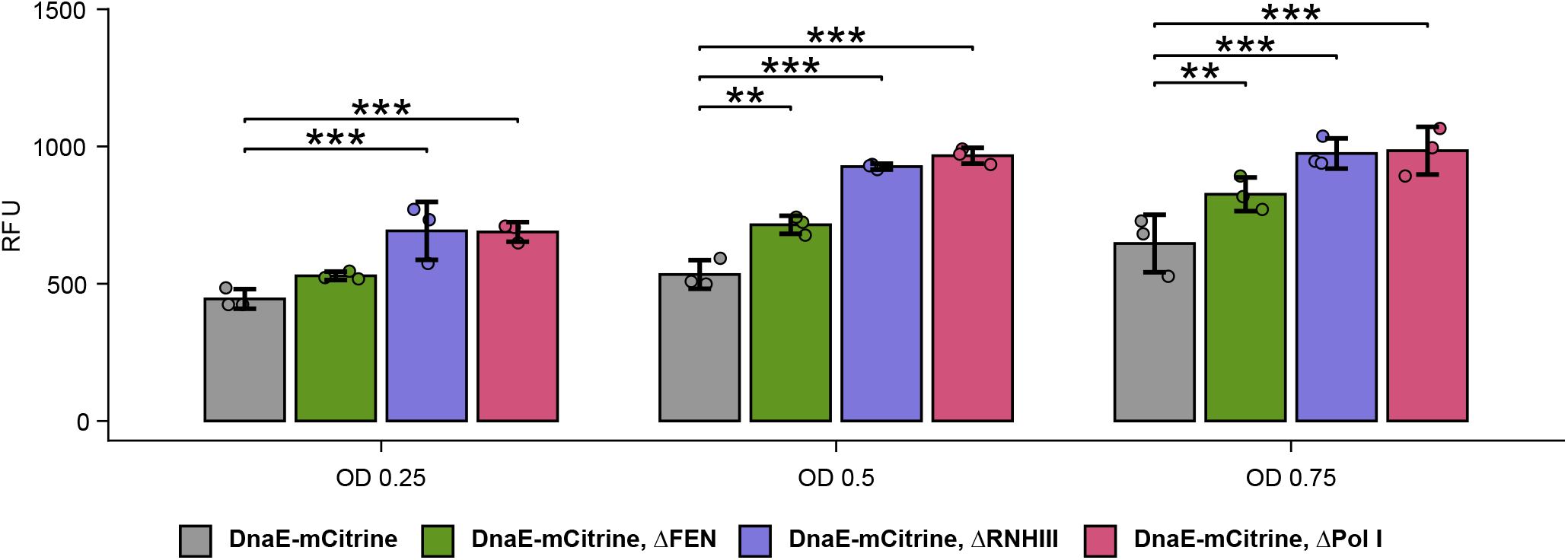
DnaE expression increases in the absence of FEN, RNase HIII, and Pol I. Visualization of results from DnaE-mCitrine fluorescence reporter assay. OD represents culture density at time of measurement. Quantification was performed for three independent replicates, with the average of two technical replicates accounted for using a linear mixed effects model. All data points are shown with ±1 SD. RFU represents fluorescent units for DnaE-mCitrine emission.

## DISCUSSION

In this work, we demonstrate two novel mechanisms of Okazaki fragment repair in *B. subtilis*. Before this study, Pol I was considered the main protein responsible for Okazaki fragment repair, using both its exonuclease domain and strand displacement synthesis abilities to degrade an RNA primer, followed by simultaneous or subsequent DNA synthesis (3,10). However, in *B. subtilis*, Pol I can be deleted with little phenotype, leading us to investigate the potential for other DNA polymerases to participate in Okazaki fragment processing. To address this question, we used purified replicative polymerases PolC and DnaE and assayed these enzymes with substrates modeled after Okazaki fragments. Using this system, we found that DnaE performs robust strand displacement synthesis from a gap, creating an RNA flap while the synthesis of DNA occurs to yield a full-length extension product. The RNA flap created by DnaE represents a substrate that can be cleaved effectively by FEN using its flap endonuclease activity (25). This mechanism accounts for both the polymerase and nuclease activity required for Okazaki fragment repair in vivo and likely represents the major repair pathway in the absence of Pol I. We also found that PolC can participate in Okazaki fragment repair when coupled with a nuclease capable of digesting the downstream ribonucleotides or a downstream flap. This mechanism likely represents a minor maturation pathway that can occur in the absence of Pol I, and it is unlikely to occur often given the robust strand displacement and DNA synthesis activity we observed by DnaE. Our work also suggests that since PolC cannot displace an annealed 5′ end, this provides a mechanism to halt DNA synthesis to allow for adjacent Okazaki fragments to be joined by ligase following FEN activity.

Overall, the findings of this study reveal two novel mechanisms of Okazaki fragment repair in *B. subtilis*, both of which occur without Pol I. In conjunction with other work (23,25), the data shown here lead us to propose four potential pathways of Okazaki fragment maturation in *B. subtilis* **(Figure 6)**. The canonical *E. coli* pathway involves RNA primer incision by RNase HIII, followed by primer removal and DNA synthesis by Pol I (23,25). The FEN pathway, typified by *B. subtilis*, involves primer degradation by FEN and synthesis by Pol I, and likely represents the major Okazaki fragment repair pathway in the presence of Pol I for organisms with an active secondary FEN (25). In the absence of Pol I, we show that the DnaE pathway mediates the majority of repair, displacing the primer using strand displacement synthesis. This can then be followed by RNA primer removal by FEN. Moreover, should FEN excise the RNA primer before the polymerase reaches the downstream Okazaki fragment, PolC can perform DNA synthesis to yield a full-length product.

**Figure 6.**
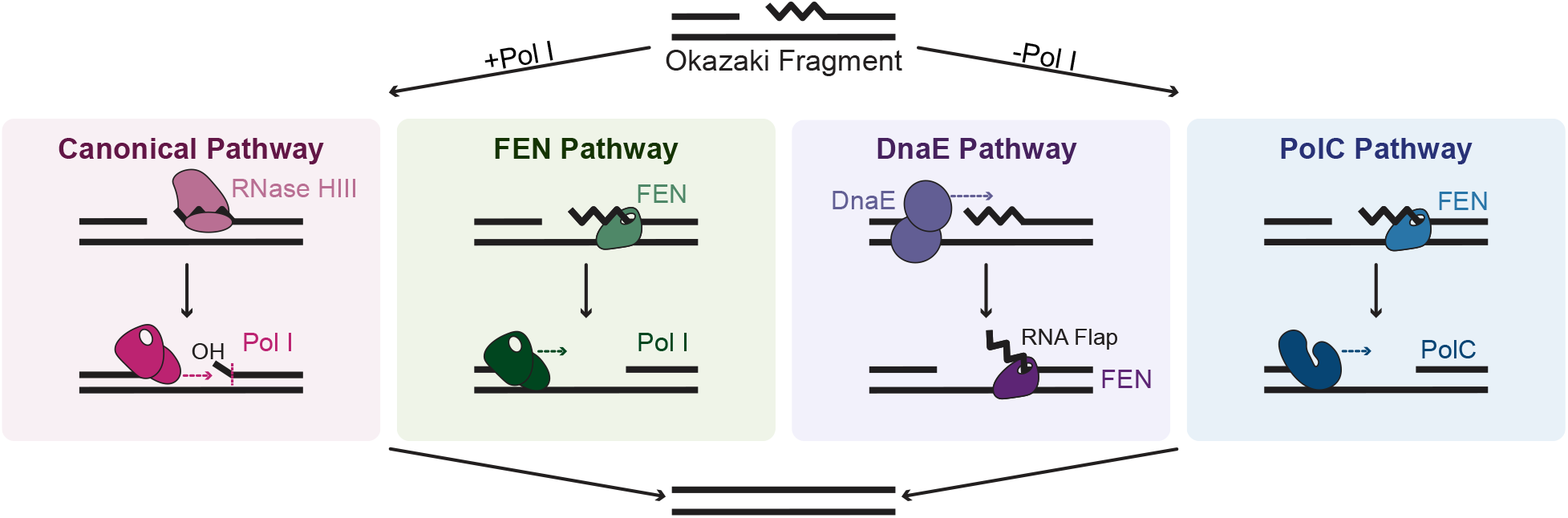
Model of four proposed pathways of Okazaki fragment repair in *B. subtilis*. The canonical pathway in most bacteria involves incision by RNase HIII/HI, leaving a few ribonucleotides that can be excised by Pol I, followed by Pol I-mediated DNA synthesis. The FEN pathway involves complete primer removal by FEN and DNA synthesis by Pol I, and represents the major repair pathway in *B. subtilis*. Canonical and FEN pathways only occur in the presence of Pol I. The DnaE proposed pathway involves strand displacement by DnaE, followed by primer excision by FEN. The PolC proposed pathway involves RNA primer degradation by FEN, followed by DNA synthesis by PolC. The DnaE and PolC proposed pathways occur in the absence of Pol I.

This work uncovers previously unknown similarities between bacterial and eukaryotic Okazaki fragment repair. In eukaryotes, DNA polymerase α synthesizes an RNA primer using its primase subunits before switching to DNA synthesis. This DNA polymerase has poor processivity and hands off the substrate to DNA polymerase δ which completes the synthesis of the fragment (5). In *B. subtilis*, primase (DnaG) synthesizes RNA, which is extended by DnaE. Thus, these two proteins effectively combine to fulfill the same role as eukaryotic Pol α (7). After DnaE extends the RNA primer, PolC completes synthesis, analogous to eukaryotic Pol δ (45). Additionally, Pol δ can perform strand displacement, leading to the formation of an RNA primer flap that becomes a cleavage substrate for FEN1 (45). Although we show that PolC is poor at strand displacement synthesis activity, DnaE can extend an upstream fragment of DNA while displacing an RNA primer, creating a substrate for FEN cleavage and synthesizing a full-length strand of DNA. Our DnaE-proposed mechanism identifies both a polymerase and nuclease (FEN) for Okazaki fragment repair (25) that can operate independently of Pol I, describing a system that bears similarity to the well-characterized eukaryotic repair mechanism (45).

Moreover, we show that the strand displacement activity of DnaE is not limited to an RNA-DNA hybrid substrate. We show that DnaE can perform strand displacement on a DNA-only substrate, suggesting that DnaE may be involved in broader DNA repair mechanisms beyond Okazaki fragment maturation. This supports previous findings, which suggest that the active site of *B. subtilis* DnaE is relatively relaxed and thus able to bypass lesions such as abasic sites or AAF adducts, unlike other replicative polymerases (44). The lesion bypass ability of *Bs*DnaE was also more robust than that of *Ec*DnaE (43,44). However, these authors found that overproduction of *Bs*DnaE does not result in spontaneous mutagenesis, suggesting DnaE is not a true error-prone polymerase (43). In our work, we find that *Bs*DnaE can perform strand displacement without an existing flap, suggesting that DnaE could act as a translesion synthesis polymerase in contexts where strand displacement synthesis is required to bypass DNA lesions.

In addition to its essential role as a major replicative polymerase, our work demonstrates that DnaE can participate in Okazaki fragment repair using its strand displacement synthesis activity. As such, DnaE could represent a potential antimicrobial drug target, along with other replicative polymerases. Since replication is essential for cell viability, various components of the replisome are potential targets for therapeutic interventions (46). While no major replicative polymerase inhibitors have reached clinical development, three main classes of drugs are known to inhibit the α subunit of DNA polymerase III: 6-anilinouracils (AUs), guanine inhibitors, and non-nucleobase inhibitors (47). Most synthesized compounds in this class target PolC in gram-positive bacteria and compete with the DNA template or nucleotides (47). The most well-established drug is HPUra, a dGTP analog, that is currently used in lab settings (48). Several drugs have been modeled after this specific compound, with the goal of limiting primer elongation through competition with nucleotides (49-52). These drugs could be effective therapeutics since C-family polymerases are exclusively found in bacteria, and thus an inhibitor could avoid cytotoxicity by sparing eukaryotic (host) polymerases (53). C-family polymerases are highly conserved throughout bacteria, although they differ in terms of lineage, function, and structure, and they have been divided into four distinct groups by phylogenetic analysis: PolC, DnaE1, DnaE2, DnaE3 (53). As a result, the diversity of this polymerase family could allow for a variety of drug development opportunities with specificity to bacterial targets. Taken together, our work highlights the previously unrecognized complexity of Okazaki fragment repair in *B. subtilis* and underscores the importance of revisiting fundamental molecular processes in bacteria.

## Author Contributions

A.H.K., F.C.L., L.A.S: conceptualization, A.H.K, F.C.L., L.A.S., L.G.J.,: investigation, A.H.K., F.C.L, L.A.S: formal analysis, A.H.K., F.C.L, L.A.S: writing-review and editing.

## ACKNOWLEDGEMENTS

We thank members of the Simmons lab for their helpful contributions during the development of this work. This work was funded by National Institutes of Health grant R35GM131772 to LAS. We thank Charles McHenry for purified PolC and DnaE proteins.

