## Supplementary material for "DnaE uses strand displacement synthesis during Okazaki fragment repair": Figures and Tables

Short title: DnaE functions during Okazaki fragment repair

**Keywords:** *Bacillus subtilis,* DNA replication, Okazaki fragments, FEN, RNase HIII

**SUPPLEMENTAL INFORMATION**

**Table S1. Strains used in this study.**

| Strain | Background | Genotype | Source |
|---|---|---|---|
| AHK27 | PY79 | Spβ$^0$ prototroph | (1) |
| FCL15 | PY79 | *dinC-gfp* (spc$^r$) | (2) |
| FCL16 | PY79 | Δ*rnhC, dinC-gfp* (spc$^r$) | This work |
| FCL9 | PY79 | Δ*rnhC,* Δ*fenA, dinC-gfp* (spc$^r$) | This work |
| FCL4 | PY79 | Δ*fenA* | (3) |
| FCL11 | PY79 | Δ*rnhC* | (3) |
| FCL3 | PY79 | Δ*polA* | (3) |
| FCL5 | PY79 | Δ*fenA,* Δ*rnhC* | (3) |
| FCL12 | PY79 | Δ*rnhC, polA* | (3) |
| AHK85 | PY79 | *dnaE::dnaE-mCitrine* | JWS163 (lab stock) |
| AHK114 | PY79 | *dnaE::dnaE-mCitrine,* Δ*polA* | This work |
| AHK112 | PY79 | *dnaE::dnaE-mCitrine,* Δ*fenA* | This work |
| AHK113 | PY79 | *dnaE::dnaE-mCitrine,* Δ*rnhC* | This work |
| AHK125 | PY79 | *dnaE::dnaE-mCitrine, dnaX::dnaX-mCherry* | This work |
| AHK128 | PY79 | *dnaE::dnaE-mCitrine, dnaX::dnaX-mCherry,* Δ*polA* | This work |

All strains are derivative of PY79.

**Table S2. Oligonucleotides used in this study**

| Oligonucleotide | Purpose | Sequence (5′-3′) |
|---|---|---|
| oFCL6 | Template | GCAATCGACTCGTAAGCATGGTTCACTACTAGCTGCACATCGCTGCTTGATGCTCAATCG |
| oFCL8 | Primer | /5IRD800/C*G*A*TTGAGCATCAAGCAGCG |
| oFCL11 | Ladder | /5IRD800/CGATTGAGCATCAAGCAGCGATGTGCAGCTAGTAGTGAACCATGCTTACGAGTCGA TTGC |
| oFCL28 | Template | GCAATCGACTCGTAAGCAGTTGGACAGCAGAGCTGCACATCGCTGCTTGATGCTCAATCG |
| oFCL30 | Downstream fragment | AGTAGTGAACCATGCTTACGAGTCGATTGC/3IR800CWN/ |
| oFCL31 | Primer | /5IRD700/C*G*ATTGAGCATCAAGCAGCG |
| oFCL32 | Ladder | /5IRD700/CGATTGAGCATCAAGCAGCGATGTGCAGCTAGTAGTGAACCATGCTTACGAGTCGA TTGC |
| oJR361 | Template | GCAATCGACTCGTAAGCATGGTTCACTACTCGCTGCTTGATGCTCAATCG |
| oJR362 | Primer | /5IRD800/CGATTGAGCATCAAGCAGCG |
| oJR367 | Downstream fragment | rArGrUrArGrUrGrArArCrCrATGCTTACGAGTCGATTGC/3IRD700CWN/ |

IRD indicates an infrared dye with the excitation noted (700 or 800 nM).
* indicates a phosphorothioate linkage. Ribonucleotides are denoted by a lowercase 'r'.

**Table S3. Substrates used in this study**

| Substrate Type | Oligonucleotides | Ladder |
|---|---|---|
| Primed | oJR361, oJR362 | oJR362, oFCL11 |
| Nicked | oJR361, oJR362, oJR367 | oJR362, oFCL11 |
| 10 nt gap | oFCL6, oFCL8, oJR367 | oFCL8, oFCL11 |
| 10 nt gap with flap | oFCL28, oFCL8, oJR367 | oFCL8, oFCL11 |
| 10 nt gap (DNA only) | oFCL6, oFCL31, oFCL30 | oFCL31, oFCL32 |

**Table S4. Plasmids used in this study**

| Plasmid Identifier | Vector | Insert |
|---|---|---|
| pFCL1 | his-pE-SUMO | *fenA* (3) |
| pFCL3 | his-pE-SUMO | *polA* (3) |
| pFCL22 | his-pE-SUMO | *rnhC* |

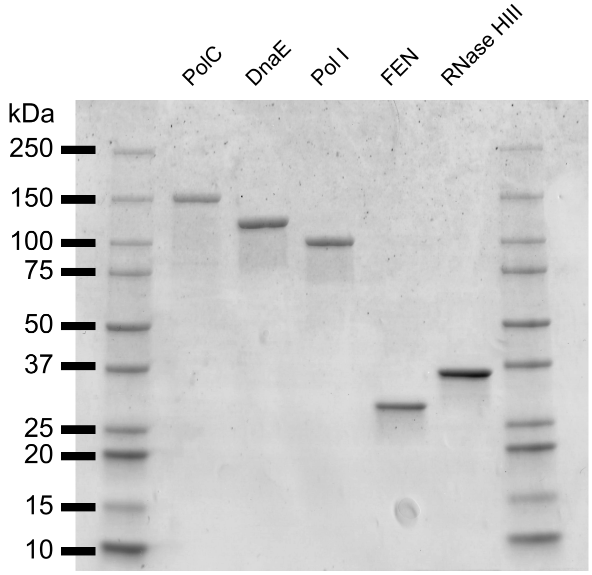


**Figure S1. Proteins used in extension assays are clean purifications.**
A total of 2 µg of each protein was separated following purification on SDS-PAGE as detailed in the Materials and Methods. SDS-PAGE was visualized following staining with Coomassie blue.

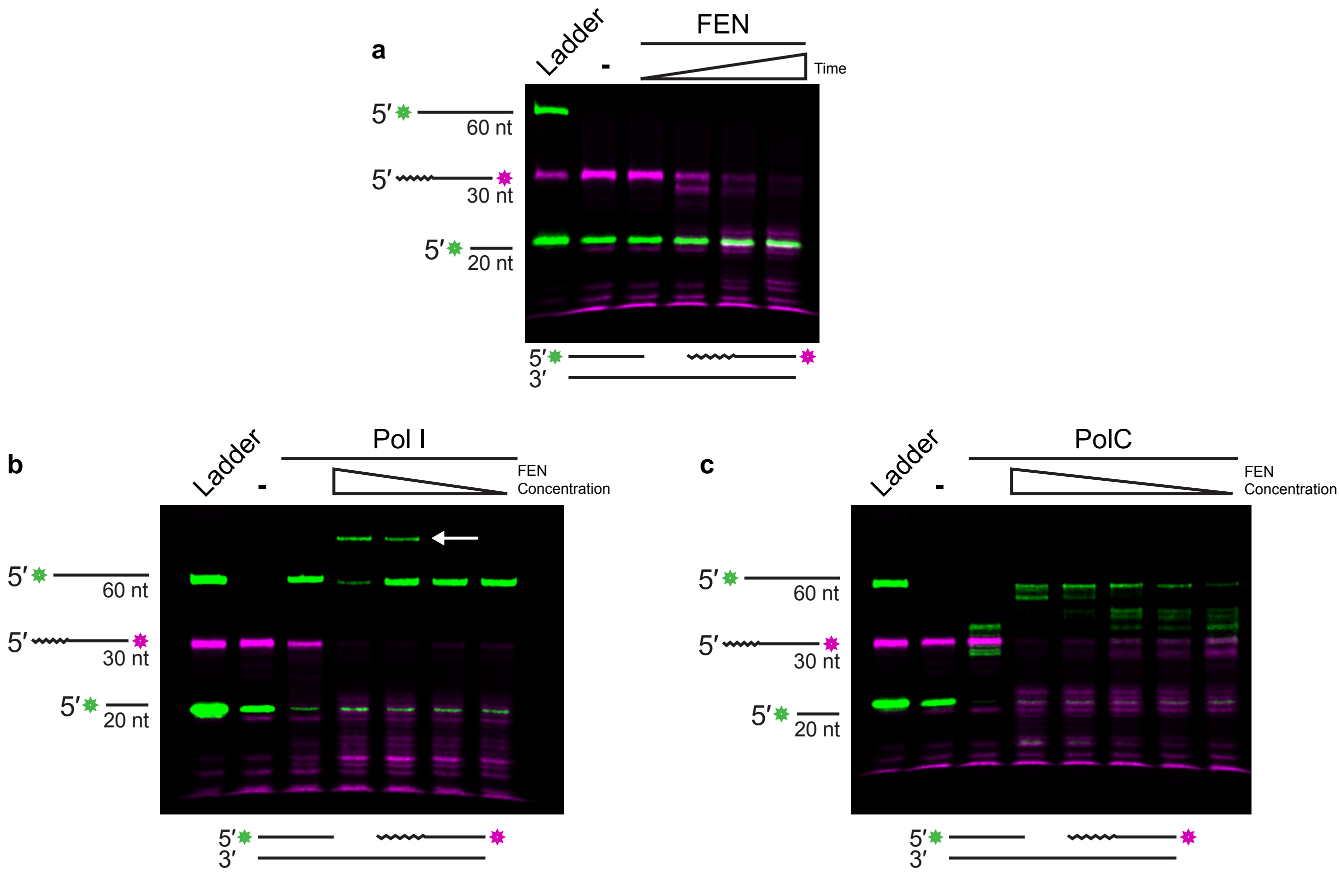


**Figure S2. Relative degradation and extension products depend on protein concentrations.**
**(A)** Products of an extension assay using a 10 nt gap substrate visualized by urea-PAGE. Ladder shows length of primer, downstream fragment, and full-length extension product. Embedded ribonucleotides denoted by squiggly line. Representative time increases from left to right with the following time points: 0 min, 1 min, 5 min, and 10 min.
**(B)** Products of an extension assay using a 10 nt gap substrate visualized by urea-PAGE. Ladder shows length of primer, downstream fragment, and full-length extension product. Embedded ribonucleotides denoted by squiggly line. FEN concentration decreases from left to right with the following concentrations: 50 nM, 25 nM, 10 nM, and 5 nM. Pol I concentration is consistent in all lanes (100 nM), except no protein control. Gel shift is denoted by white arrow.
**(C)** Products of an extension assay using a 10 nt gap substrate visualized by urea-PAGE. Ladder shows length of primer, downstream fragment, and full-length extension product. Embedded ribonucleotides denoted by squiggly line. FEN concentration decreases from left to right with the following concentrations: 50 nM, 25 nM, 15 nM, 10 nM, and 5 nM. PolC concentration is consistent in all lanes (100 nM), except the no protein control.
